# Sniff-invariant intensity perception using olfactory bulb coding of inhalation dynamics

**DOI:** 10.1101/226969

**Authors:** Rebecca Jordan, Mihaly Kollo, Andreas T Schaefer

**Author notes:** Correspondence and requests for materials, including custom software, should be addressed to A.T.S via email to.

## Abstract

For stable perception of odor intensity, there must exist a neural correlate that is invariant across other parameters, such as the highly variable sniff cycle. Previous hypotheses suggest that variance in inhalation dynamics alters odor concentration profiles in the naris despite a constant environmental concentration. Using whole cell recordings in the olfactory bulb of awake mice, we directly demonstrate that rapid sniffing mimics the effect of odor concentration increase at the level of both mitral and tufted cell (MTC) firing rate responses and temporal responses. In contrast, we find that mice are capable of discriminating concentrations within short timescales despite highly variable sniffing behavior. We reason that the only way the olfactory system can differentiate between a change in sniffing and a change in concentration is to receive information about the inhalation parameters in parallel with information about the odor. While conceivably this could be achieved via corollary discharge from respiration control centres, we find that the sniff-driven activity of MTCs without odor input is informative of the kind of inhalation that just occurred, allowing rapid detection of a change in inhalation. Thus, a possible reason for sniff modulation of the early olfactory system may be to inform downstream centres of nasal flow dynamics, so that an inference can be made about environmental concentration independent of sniff variance.

---

For consistency of perception, sensory systems must be able to stably encode the same perceptual features across a wide range of situations. An example of this is the encoding of object size independent of object distance in the visual system (Helmholtz, 1867) – we do not perceive a giant apple when viewed at close range, and similarly we do not misperceive buildings as tiny objects when viewed from great distance. In the olfactory system, studies have looked at how odor identity may be encoded independent of odor concentration (Cleland et al., 2012; Uchida and Mainen, 2008; Wachowiak et al., 2002; Wilson et al., 2017) and sniff cycle variance (Cury and Uchida, 2010). An olfactory problem that has received less attention, however, is stable encoding of odor intensity – the perceptual correlate of odor concentration (Wojcik and Sirotin, 2014). Increasing concentration is known to affect neural activity in many ways (Mainland et al., 2014). At the level of glomerular input from olfactory sensory neurons (OSNs), increasing concentration enhances the activity of already responsive glomeruli and incorporates new glomeruli into the activity profile, overall resulting in a broadening of the spatial ‘map’ of activity (Rubin and Katz, 1999; Spors and Grinvald, 2002). Changes in spike rate are also seen at the level of the olfactory bulb output cells, mitral and tufted cells (MTCs), though this can be a more complex mixture of inhibitory and excitatory effects (Bathellier et al., 2008; Cury and Uchida, 2010; Fukunaga et al., 2012; Meredith, 1986), and is thought to be constrained via inhibitory circuits (Kato et al., 2013; Miyamichi et al., 2013; Roland et al., 2016). The perhaps more ubiquitous correlates of concentration increase, however, are temporal response changes, notably with early excitation undergoing a latency reduction in OSNs (Ghatpande and Reisert, 2011; Rospars et al., 2000), MTCs (Cang and Isaacson, 2003; Fukunaga et al., 2012; Sirotin et al., 2015), as well as in the piriform cortex (Bolding and Franks, 2017). This is thought to arise since OSNs will depolarise to threshold more quickly when the concentration profile in the naris is steeper.

In awake mice, sniffing behaviour is in continual flux (Kepecs et al., 2007; Welker, 1964; Wesson et al., 2008a, 2009; Youngentob et al., 1987). This might present a problem for concentration coding: firstly, changing flow will affect the number of odor molecules entering the nasal passage, altering the concentration profile in the naris despite a stable environmental concentration (Mainland and Sobel, 2006; Shusterman et al., 2017; Teghtsoonian et al., 1978). Secondly, altering the velocity of air in the naris will alter the time at which odorised air reaches the olfactory epithelium, which may disrupt the reliability of temporal coding of concentration (Shusterman et al., 2017). Despite this, previous work suggests that humans can perceive odor intensity independent of the inhalation flow rate (Teghtsoonian et al., 1978). Here, using whole cell patch recordings in awake mice, we show that faster sniffs evoke response changes identical to those caused by increasing concentration. Surprisingly however, we show that variance in sniffing has little effect on the performance of mice trained to make fine concentration discriminations. Finally, we propose that the olfactory system can make an inference about whether a response change was caused by concentration change or sniff change by encoding the parameters of sniffing on fast timescales in mitral and tufted cells, which respond to inhalation change in cell-type specific ways, allowing rapid detection of a change in sniffing.

## Results

### Changes in sniffing can mimic the effect of increased concentration on firing rate response

We first wanted to determine whether the effect of sniff changes on MTC odor response could qualitatively mimic concentration changes at the level of FR change. To do this, we used whole cell recordings from identified MTCs in awake passive mice, as this allows unbiased sampling from the MTC population in terms of baseline firing rate (FR), and reliable identification of cell type based on electrophysiology (Kollo et al., 2014; Margrie et al., 2002). On each trial, mice were presented randomly with 2 s long odor stimuli calibrated to either 1% (low concentration) or 2.5% (high concentration) square pulses. On a small percentage of low concentration trials, mice also received a gentle air puff to the flank, evoking fast sniffing behavior characterised by high frequency sniffs and short inhalation duration (Fig. 1A and Supplementary Fig. 1). For all analyses in the manuscript, ‘odor onset’ (t=0) is defined as the first inhalation onset during the odor stimulus. Note that several parameters of sniffing co-vary with inhalation duration, including the sniff duration, the previous sniff duration and the slope of the inhalation (Supplementary Fig. 2). Thus wherever we refer to ‘fast’ or ‘slow’ sniffing, this will necessarily refer to variance in these multiple parameters.

**Figure 1.**
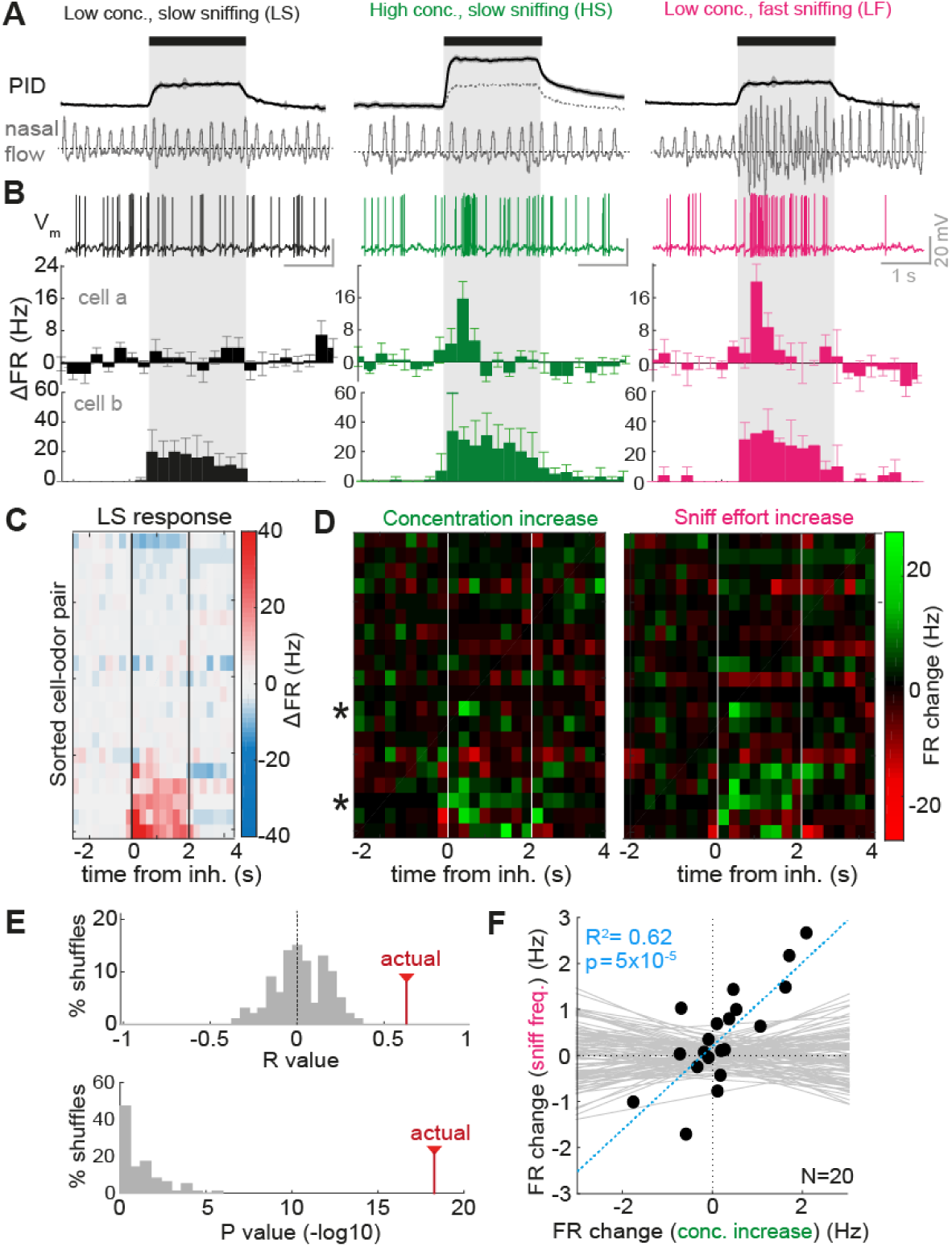
Sniff change and concentration change have similar effects on FR responses of MTCs. (**A**) Stimulation paradigm during whole cell recordings. PID traces show response of photoionisation detector (magnitude proportional to odor concentration), while nasal flow traces show example sniffing recorded using external flow sensor for the three types of trial. Black bar and grey box shows where odor is on, aligned to first inhalation onset. (**B**) Example odor responses recorded in each stimulus condition. V_m_ traces show example responses for cell a, while PSTHs below show averaged FR responses in 250 ms time bins for 5 trials in each case. Bottom-most PSTHs are calculated for a different example, cell b. Error bars show standard deviation. All are aligned to first inhalation onset. (**C**) Heatmap of average FR respones for all cell odor pairs in the low concentration slow sniff frequency condition (LS), ordered by mean FR response. (**D**) Heatmap of FR response differences (difference between PSTHs): Concentration increase = HS-LS, sniff frequency increase = LF-LS. Asterisks indicate cell a and cell b examples. (**E**) Top: R values for correlation across all odor time bins as shown in D, between FR change due to concentration change and those due to sniff frequency change. Histogram shows R values for shuffle controls, ‘actual’ shows R value for real data. Bottom: as for above, but histogram showing p-values for the correlations (-log_10_) (**F**) Correlation between mean FR response change for concentration change (HS-LS) and sniff frequency change (LF-LS) across first second of odor stimulus. N = 20 cell-odor pairs. Grey lines show correlations calculated for 100 shuffle controls, blue line shows real correlation.

During recordings, it was apparent that some cells displayed overt changes in FR with the increase in concentration, and the most salient of these were increases in excitatory FR response (Fig. 1B, cell a and cell b). When comparing changes in FR evoked by concentration increase to those taking place as a result of increased sniff frequency, it was apparent that very similar changes took place (Fig. 1B). Altogether we recorded from 20 MTC pairs in such a manner, with a range of FR responses to the low concentration odorant (Fig. 1C). Comparing heat maps of the changes in FR due to increased concentration and due to increased sniff frequency revealed a very similar set of changes across the dataset (Fig. 1D), which were significantly correlated compared to shuffle controls (R = 0.63, p = 5×10^−19^; Fig. 1E; see methods). When taking a broad measure of the change in firing rate across the first second of the stimulus, changes in FR were significantly correlated between those resulting from concentration increase and those resulting from faster sniffing (R^2^ = 0.62, p = 5×10^−5^, n = 20; Fig. 1F).

While in the output of MTCs the effect of sniffing and concentration increase were very similar, differences were seen in the subthreshold response changes, suggesting that changes in input in the two cases were not identical: increases in inhibition were generally larger for concentration increase than for faster sniffing (Supplementary Fig. 3). We suggest this could be the result of inhibitory networks which act to normalise olfactory bulb output (within limits) in the face of increased input (Kato et al., 2013; Miyamichi et al., 2013; Roland et al., 2016).

Thus, while increased concentration causes greater increases in subthreshold inhibition than increased sniff frequency, the latter results in changes in the olfactory bulb output that apparently mimic those resulting from increases in concentration.

### Faster inhalation causes temporal shifts similar to those reported for concentration increase

It has been reported that increased concentration causes changes in response on finer temporal timescales, in particular the advance of excitatory bursts (Cang and Isaacson, 2003; Fukunaga et al., 2012; Sirotin et al., 2015). Does a faster sniff similarly cause such temporal shifts on early timescales?

To determine this, we first analysed 13 cell-odor pairs with early excitatory responses recorded in passive mice where only a single concentration stimulus (1% saturated vapour pressure) was presented to the animal across trials. Comparing the FR response over the first 250 ms for ‘fast’ sniff trials (>70^th^ percentile peak inhalation slopes), and ‘slow’ sniff trials (<30^th^ percentile), it was apparent that faster inhalation could cause a latency advance of the excitatory burst (Fig. 2A–B). Across all cell-odor pairs, faster inhalation caused a significant latency reduction in mean response onset (latency change (fast-slow) = −16 ± 14 ms, p = 0.002 paired t-test between onsets for slow and fast inhalations; Fig. 2C). Onset latencies displayed a significant relationship with the peak firing rate during the response (R^2^ = 0.61, p = 0.002, n = 13; Fig. 2E), suggesting that the most strongly activated cells respond earlier. The extent of the latency reduction for fast sniffing was significantly correlated with the onset time during slow inhalation: if the response was of longer latency during slow sniffing, the latency reduction was greater (R^2^ = 0.46, p = 0.01, n = 13; Fig. 2D). This indicates that those cell-odor pairs showing a stable latency are likely already responding at the earliest timescale – i.e. there is a lower limit on the latency of odor response.

**Figure 2.**
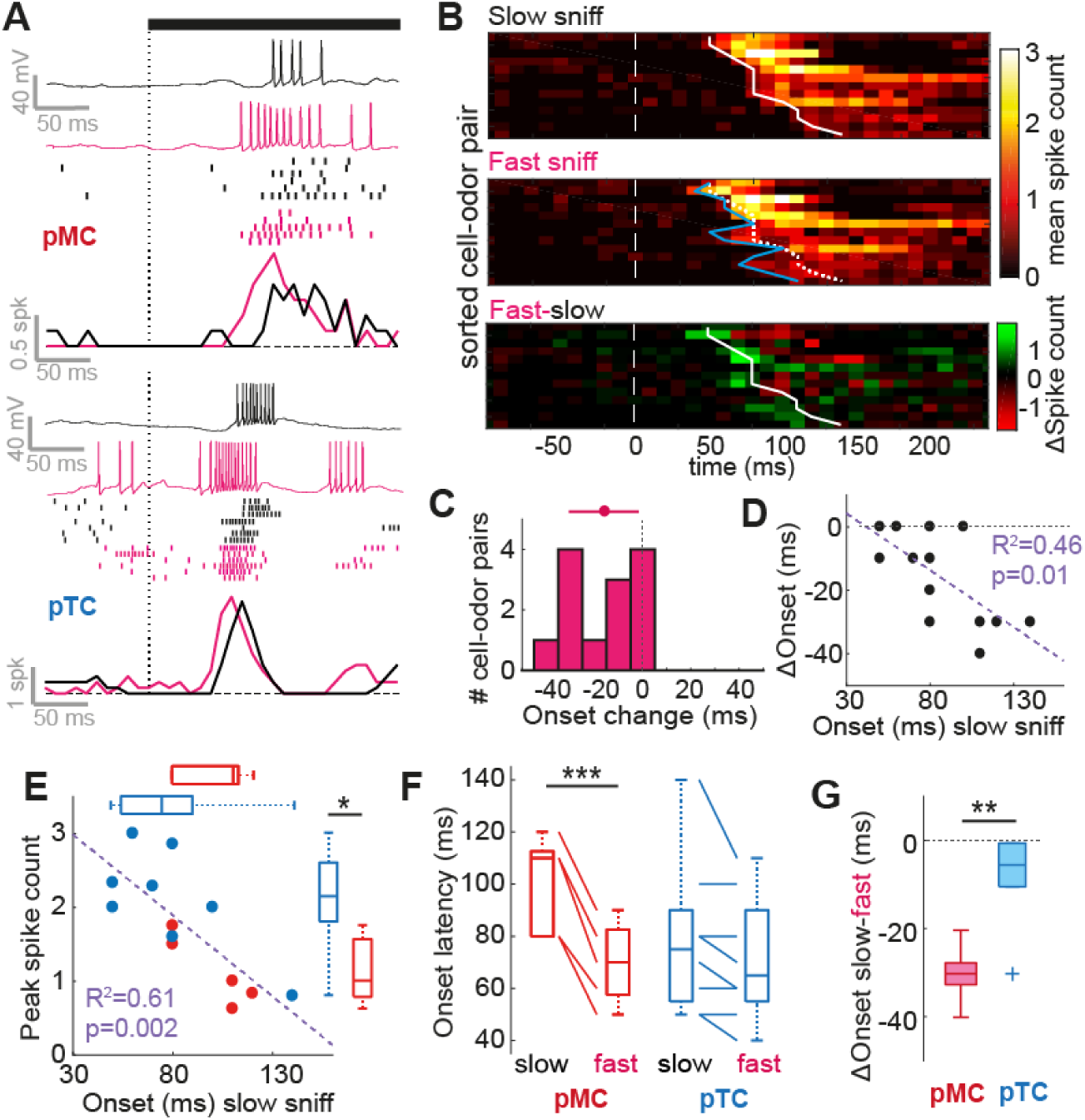
Faster inhalation causes temporal shifts similar to those reported for concentration increase. (**A**) Example V_m_ traces, spike rasters and mean spike counts for early excitatory responses for slow inhalation (black) and fast inhalation (pink), for two different cell-odor pairs. The top example is from a putative mitral cell (pMC) and bottom example is from a putative tufted cell (pTC). Rasters are ordered (top to bottom) by slowest to fastest inhalation. Black bar and dotted line indicate odor onset aligned to the first inhalation onset. (**B**) Heatmaps of mean spike count for 13 cell-odor pairs showing early excitation in response to slow inhalation (top) and fast inhalation (middle). White dashed line indicates odor onset aligned to the first inhalation onset. Cell odor pairs are sorted from short to long onset latency (during slow inhalation). Bottom heatmap shows the difference between the two above (fast-slow). White solid and dotted line indicates onset latency of each cell-odor pair for slow inhalation. Blue line indicates onset latency for fast inhalation. (**C**) Histogram of onset latency changes (fast-slow) for all 13 cell-odor pairs. (**D**) Scatter plot to show relationship between onset latency for slow inhalation, and the onset change between fast and slow inhalation. (**E**) Correlation between response onset latency and peak spike count for early excitatory odor responses evoked by a slow sniff. Blue data comes from pTCs and red data comes from pMCs. Boxplots compare the two parameters for pTCs and pMCs. (**F**) Comparison of response onset latencies for excitatory responses evoked by fast and slow sniffs for pMCs and pTCs. (**G**) Comparison of response onset latency change (fast-slow sniff) for pMCs and pTCs.

It was previously shown in anaesthetized mice that mitral cells (MCs) undergo robust reductions in latency of excitation for concentration increase, while tufted cells (TCs) – which already respond earlier - do not (Fukunaga et al., 2012). We used sniff cycle phase preference boundaries to determine putative MC (pMC) and TC (pTC) phenotype using subthreshold activity as previously described (Fukunaga et al., 2012; Jordan et al., 2017). Examples could be found where both pMCs and pTCs underwent reductions in latency of excitation when the sniff was fast (Fig. 2A), however in general, reductions for pMCs were greater than reductions for pTCs (pMCs: latency change = −30 ± 7 ms, p = 7×10^−4^, paired t-test, n = 5 cell-odor pairs; pTCs: latency change = −8±10 ms, p = 0.08, paired t-test, n = 8 cell odor pairs; pMCs vs pTCs: p = 0.001, unpaired t-test; Fig. 2F–G), and this was largely because pTCs already tended to respond with shorter latency during slow sniffs than pMCs (pTC onset median = 75 ms, IQR = 55-90 ms; pMC onset median = 110 ms, IQR = 80-113 ms, p = 0.13, Ranksum; Fig. 2E).

Thus, the temporal shifts and cell-type specificity in the effect of faster sniffing matches that previously described for concentration increases in anaesthetized mice (Fukunaga et al., 2012).

### Faster inhalation mimics effect of concentration increase on latency response in the timescale of a single sniff

Could the effect of sniffing on latency directly mimic concentration effects at this timescale? When comparing high and low concentration stimuli over the first 250 ms in MTC recordings from passive mice (dataset as in Fig. 1), the only salient changes in response to increased concentration were latency advances of excitatory burst stimuli (Fig. 3A and B). When correlating changes in spike count as before (Fig. 1E) between those occurring for sniff change and those occurring for concentration change, there was a highly significant positive correlation between the two (R = 0.71, p = 4×10^−72^, n =525 time bins; Fig. 3C). Latency reductions for concentration increase were similar in magnitude to those seen due to sniff change (Fig. 3D, mean onset advance = −18±10ms, p = 0.04, n = 4 paired t-test between onsets for low and high concentration), and similar to those previously reported (Sirotin et al., 2015). This latency change contributed to the entirety of the discriminability of the two different concentrations on this timescale, with the Euclidean distance between the two dropping to baseline if the excitatory bursts were manually shifted forward for the low concentration (Fig. 3E, ‘slow sniff’ vs ‘slow sniff adv.’, see methods). Faster inhalations during low concentration trials mimicked the latency shifts caused by concentration increase, rendering the high and low concentration stimuli indistinguishable (Fig. 3E, ‘slow sniff’ vs ‘fast sniff). Registering spikes to sniff cycle phase revealed a potential increase in the ability to detect changes in concentration despite changes in sniffing (Supplementary Fig. 4; supplementary information), suggesting that the OB may require information about sniff parameters in order to determine the concentration. Thus, even on short timescales, a more rapid inhalation mimics concentration increases at the level of the olfactory bulb output.

**Figure 3.**
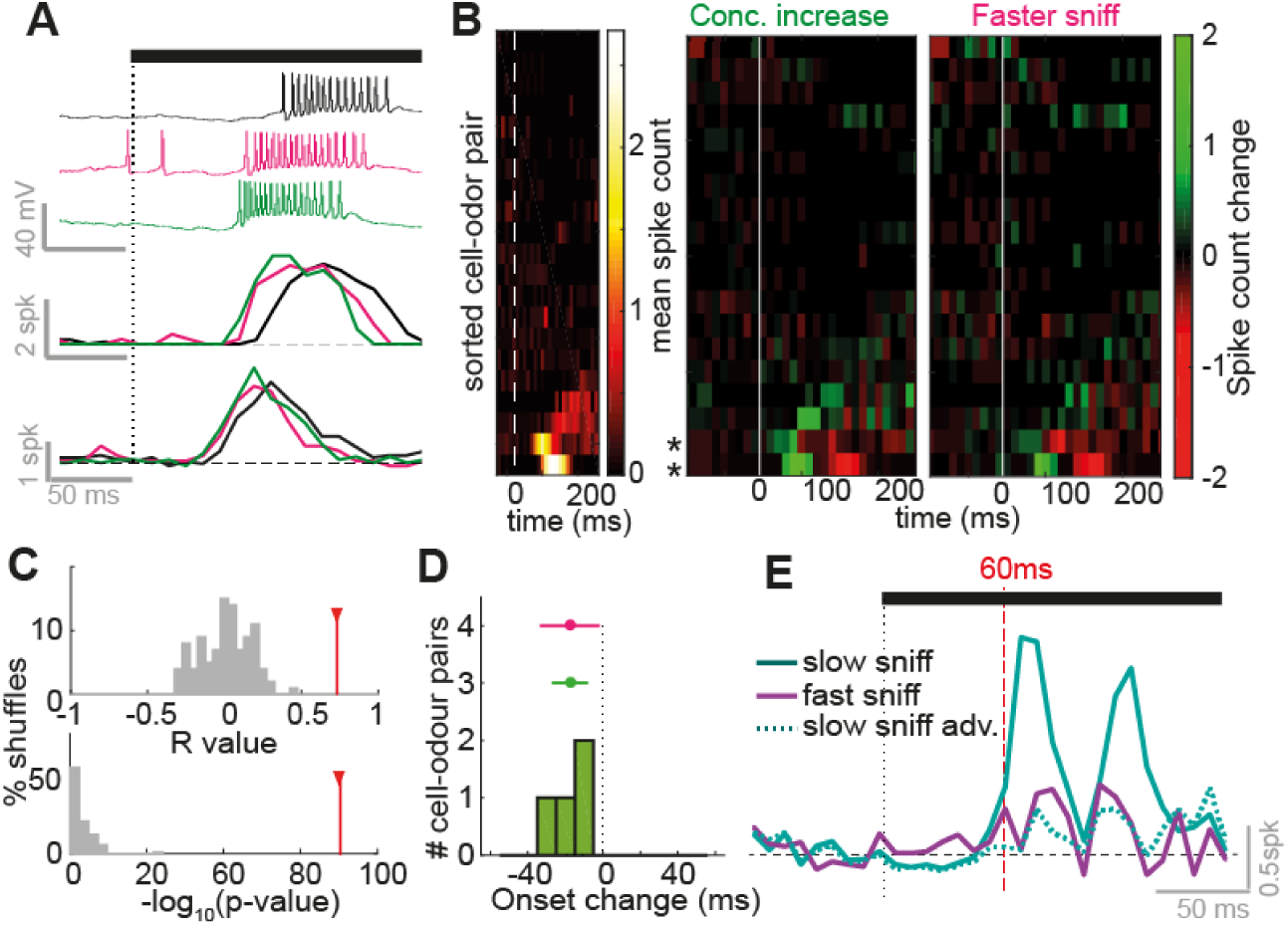
Inhalation change and concentration change cause similar temporal effects on early responses. (**A**) Example traces and mean spike counts for early excitatory responses. Black shows response at low concentration evoked by slow inhalation, pink shows response at low concentration evoked by fast inhalation, and green shows response for high concentration evoked by slow inhalation. (**B**) Left: Heatmap to show early spike counts of all 20 cell-odor pairs recorded for low concentration and slow inhalation. Cell-odor pairs are sorted by the mean spike count during odor, from low to high. Middle: Heatmap to show difference in spike counts between high concentration and low concentration (evoked by slow inhalation). Left: heatmap to show difference in spike count between fast inhalation and slow inhalation (low concentration stimulus). (**C**) Top: R values for correlation across all odor time bins as shown in B, between spike count change due to concentration increase and due to faster inhalation. Histogram shows R values for shuffle controls, red bars show R value for real data. Bottom: as for above, but histogram showing p-values for the correlations (-log10). (**D**) Histogram to show response onset changes due to concentration increase. Errorbar in green shows mean and SD of this data, and in pink shows the distribution of changes as in panel C for comparison. (**E**) Euclidean distance between population spike count response vectors for high and low concentration (where data came from slow inhalation trials; ‘slow sniff’, solid cyan), for high concentration and time-shifted low concentration (‘slow sniff adv.’; where excitatory latency changes due to concentration change were undone via time shifting of the data; dotted cyan), and for high concentration and low concentration where low concentration data came from fast inhalation trials (‘fast sniff’; solid purple).

### Mice can successfully discriminate concentrations on rapid timescales

Rodents have previously demonstrated the ability to discriminate odor concentrations (Abraham et al., 2004; Parthasarathy and Bhalla, 2013; Slotnick and Ptak, 1977; Wojcik and Sirotin, 2014). Given the physiology (Fig. 1–3), and that sniff parameters are constantly varying in awake mice (Wachowiak, 2011), we next sought to determine the capabilities of mice to distinguish odor concentrations in a simple head-fixed go/no-go paradigm (Fig. 4A-C), despite variance in sniffing.

**Figure 4.**
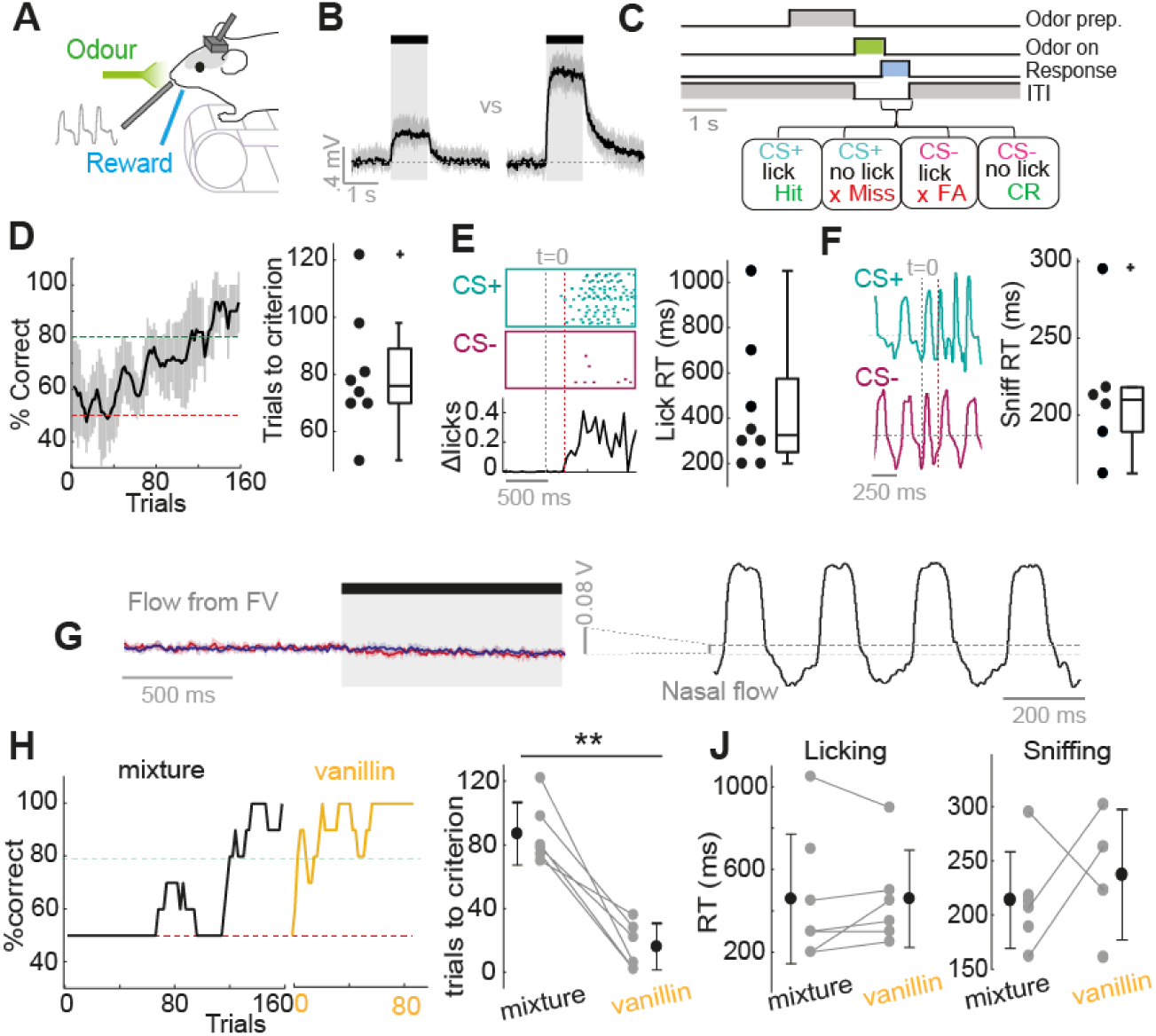
Mice rapidly learn to discriminate concentrations on fast timescales. (**A**) Diagram of head-fixed behaviour (**B**) Average PID traces for concentration go/no-go stimuli. Shaded area shows standard deviation. (**C**) Concentration go/no-go task sequence. (**D**) Left: average learning curve for 8 mice. Percentage correct is calculated as a moving average over 5 CS+ and 5 CS-trials. Shaded area indicates SD. Right: distribution of learning times to criterion (4 successive learning curve points at or above 80% correct). (**E**) Left: example to show calculation of reaction time from lick behaviour. Rasters of licking during criterion performance are shown for CS+ and CS-. A difference in mean lick count over time is then calculated (bottom) and lick reaction time (RT) is determined where this difference first exceeds 2 SDs of the baseline difference. Red dotted line indicates RT calculated for this mouse. Right: distribution of lick RTs calculated in this way. (**F**) Left: example sniff traces for CS+ and CS-during driterion performance. Prior to licking mice show rapid sniff bouts such that RT may be determined earlier using divergence of sniff waveforms (see methods). Red dotted line indicates RT calculated for this mouse. Right: distribution of RTs calculated using sniff time divergence. (**G**) Mean flow change recorded 1 mm from olfactometer output for high concentration stimulus (red) and low concentration stimulus (blue). Average of 10 trials; shaded area shows standard deviation. Y scale bar is compared to that of nasal flow traces recorded in the same manner to demonstrate the negligible nature of flow change from the olfactometer. (**H**) Learning curves for an example mouse for 2-concentration discrimination, first for the odor mixture (in black) and subsequently for vanillin (yellow). Right hand plot compares the number of trials to criterion for the initial mixture and vanillin for each mouse. (**J**) Plots to compare RTs as estimated from lick behaviour (right) and sniff behaviour (left) for the mixture stimulus and vanillin for all where a reaction time was calculable.

First, mice were trained to simply distinguish high (3%) versus low (1%) concentration stimuli. 3 mice were trained with the low concentration stimulus as the CS+ (‘Low go’), and 5 mice were trained with high concentration as the CS+ (‘High go’). After pre-training (Supplementary Fig. 5A), all mice learned the distinction within a single training session (Fig. 4D; 85 ± 19 trials to 80% correct, n = 8). Mice also made rapid decisions based on concentration: analysis of the reaction time of mice based on lick probability showed that mice could perform the task successfully within as low as 200 ms (Fig. 4E, median = 300 ms, range = 200 to 1050 ms, see methods). Similarly, using divergence of sniffing behavior between CS+ and CS-stimuli (see methods), reaction times could be estimated at 214 ± 60 ms, and as low as 160 ms (Fig. 4F) - in every case the first exhalation already showing a significant difference between CS+ and CS-stimuli. Thus mice can make decisions about concentration on the timescale of a single inhalation.

**Figure 5.**
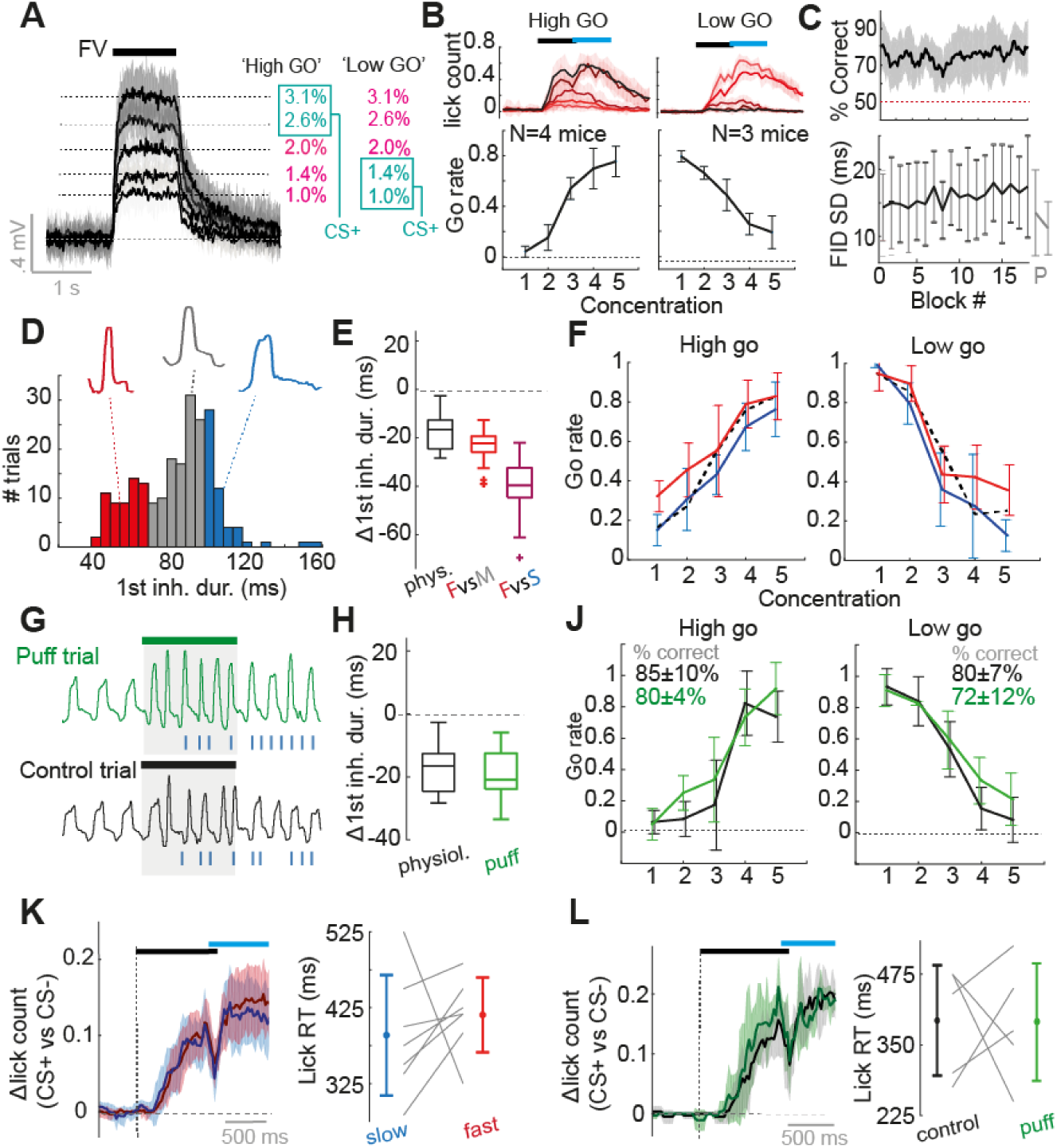
Variance in inhalation has no overt impact on concentration discrimination performance. (**A**) Diagram to show average PID traces for the 5 different concentrations and contingency schemes. Shaded area shows SD. (**B**) Top: Mean lick counts averaged across mice for the 5 different concentrations (darkest = strongest) for both ‘high go’ and ‘low go’ contingencies. Black bar indicates odor stimulus, and blue bar indicates response period. Bottom: average go rate (percentage of trials with a go response) across mice for all 5 concentrations. (**C**) Top: average % correct across all sessions of the 5 concentration discrimination task (n = 17 mouse-session pairs). Shaded area shows standard deviation. Bottom: mean standard deviation (SD) for first inhalation duration (ms) across 17 mouse-session pairs calculated for each 10-trial block of a session. Two grey points ‘P’ represent the same data but for 2 blocks in passively exposed mice (n = 26 mice). Error bars = SD. (**D**) Example histogram of inhalation durations of the first sniff during an odor stimulus across trials for one mouse. Data for each mouse is partitioned into fast inhalations (30^th^ percentile, red), slow inhalations (>70^th^ percentile, blue), and other (grey). Example representative nasal flow waveforms for a single sniff of each subset are shown. (**E**) Comparison of changes in the first inhalation between physiology and behavioural experiment. Black (‘phys.’) shows distribution of mean change used for analysis of odor responses for 20 cell-odor pairs recorded in passive mice (as in Fig. 3). Red (‘FvsM’) shows mean difference between red and grey sections of the inhalation distribution as in panel D for all mice and concentrations (n = 7 mice x 5 concentrations); Purple ‘FvsS’ shows the same as for ‘FvsM’, but shows mean difference between red and blue sections of the inhalation duration distribution as in panel D. (**F**) Go rate as a function of concentration when splitting trials according to duration of first inhalation as in D. Dotted line shows mean go rate for sniffs with inhalation between 30^th^ and 70^th^ percentile. (**G**) Example sniff traces for one animal for a puff trial (a trial in which an air puff to the flank was used to evoke fast sniffing) and an adjacent control trial of the same odor. (**H**) Comparison of changes in mean first inhalation duration for physiological analysis (Fig. 3, n = 20) and for puff vs control trials during behaviour (n = 7 mice × 5 concentrations). (**J**) Mean go rate as a function of concentration across mice for puff trials (green) vs control trials (black). (**K**) Average difference in lick-histograms between CS+ and CS-(highest vs lowest concentration) averaged across all 7 mice for slow sniff trials (blue data) and fast sniff trials (red data) partitioned as in panel D. Shaded area indicates SD. Black bar indicates odor stimulus, and blue bar indicates response period. Right plot shows difference in reaction times as measured by licking for fast and slow sniff trials for all 7 mice. (**L**) As for panel K, but now comparing lick distributions and reaction times between puff trials and control trials.

Mice were not using flow changes from the olfactometer output to make the discrimination, as our olfactometer design keeps flow from odor outlet constant (Fig. 4G). Secondly, trigeminal stimulation was likely not being used since mice subsequently learned to discriminate the same concentrations of vanillin (an odor which is thought not to stimulate trigeminal afferents – Frasnelli et al., 2011), within a significantly shorter timeframe (16 ± 14 trials to criterion, p = 0.001, paired t-test, n = 6 mice, Fig. 4H), `suggesting they may generalise the ‘task rule’ for concentrations across` different odors. Moreover, reaction times for vanillin were not significantly different than for the mixture (licking RT: p = 0.46, paired t-test, n = 6; sniffing RT: 237 ± 45 ms, p = 0.5, unpaired t-test; Fig. 4J). Finally, learning in the task was likely the result of acquiring the response to the stimulus rather than learning how to perceive the difference in concentration, since on the very first presentation of the CS-concentration after pre-training on the CS+ concentration, mice typically displayed a rapid sniffing response (Supplementary Fig. 5B) classically associated with novel odor identity (Roland et al., 2016; Verhagen et al., 2007; Wesson et al., 2008b).

Thus, mice can very rapidly make decisions based on relatively modest concentration differences within the timescale of a single sniff, comparing very well to their abilities in odor identity tasks (Resulaj and Rinberg, 2015; Uchida and Mainen, 2003; Wesson et al., 2008b).

### Variance in sniffing has no overt impact on performance in a fine concentration discrimination task

To determine whether sniff variation impacts the concentration decisions of mice, 7 trained mice were advanced on to a 5-concentration task. Here, three new intermediate concentrations between the two previously learned concentrations were presented (Fig. 5A). The concentration most similar to the learned CS+ was rewarded as a CS+, while the other two concentrations, including one exactly halfway between the previously learned concentrations, were treated as CS-(Fig. 5A). 2-3 sessions of 200 trials were performed on this task, over which mice generally performed at a high level of accuracy (Fig. 5B-C, mean percent correct across session = 75 ± 6%, n = 7 mice). Were mice learning a stable sniffing strategy to perform the task? This seems unlikely, as variance in inhalation duration of the first sniff for each mouse did not decrease over the session (if anything, a mild increase in sniff variance was observed: R = 0.72, p = 8 × 10^−4^, regression between block number and mean variance, n = 18 blocks; Fig. 5C), and variance was significantly larger compared to passive control mice (behaving median first inhalation duration SD = 0.16 ms, IQR = 12 to 20 ms, n = 284 mouse-block pairs ; passive median SD = 11 ms, IQR = 9 to 15 ms, n = 47 mouse-block pairs; p = 4 × 10^−5^, Ranksum; Fig. 5C).

Mice displayed a graded percentage of correct versus error trials across concentrations, indicating that the discrimination task was not trivial (Fig. 5B). Thus, shifts in perceived concentration should be overtly seen in the performance curves. To test this, we first separated trials according to whether the first sniff was fast (<30th percentile inhalation duration) or slow (>70^th^ percentile) (Fig. 5D). This resulted in a comparison of trials between which the difference in the inhalation duration matched or exceeded that used in the whole cell recordings when comparing fast and slow sniff trials (Fig. 5E). Recalculating performance curves for each subset, there were no large or significant differences in the performance curves for mice performing on either contingency (Fig. 5F; p>0.01 paired t-tests). Secondly, on a small selection of trials for 5 of the mice, the puff stimulus (as used during the physiological recordings) was used to evoke fast sniffs, including the first inhalation (Fig. 5G). The mean changes in first inhalation duration evoked by this puff were again highly comparable to that used for analysis of fast and slow sniffs in the physiological data (Fig. 5H). While this had a minor but insignificant effect on error rate likely owing to distraction (percent correct: control trials = 83 ± 8%, probe trials = 77 ± 9%, p = 0.16 paired t-test, n = 5 mice), there were remarkably no gross differences in the performance curves compared to control trials (p>0.01, paired t-tests; Fig. 5J). Finally, when separating trials for each concentration according to the response of the mouse (either ‘go’ or ‘no go’), there was no overt differences in first inhalation between go and no-go trials ( Supplementary Fig. 6).

Given that we have only considered the first sniff cycle, it is possible that mice take another sniff prior to making a decision if the initial sniff was fast and gave rise to ambiguity about concentration. This would be reflected in longer reaction times for fast compared to slow first sniff trials. Comparing trials with fast and slow inhalations as above (Fig. 5D), reaction times (calculated between highest and lowest concentration) were not significantly different (Δreaction time fast-slow = −30 ± 105 ms, p = 0.53, paired t-test, n = 7, Fig. 5K). Similarly, reaction times were unaffected by the puff stimulus as compared to control trials (Δreaction time probe-control = 30 ± 166 ms, p = 0.96, paired t-test, n = 5, Fig. 5L). This was also the case for finer concentration discrimination (Supplementary Fig. 6D and E). Thus, mice were not using larger amounts of information to make their decisions when sniffs were fast.

Reductions of inhalation duration of 10-20 ms rendered 1% and 2.5% concentrations indiscriminable within MTC responses (Fig. 3E-K). Here we are comparing similar and even larger reductions in inhalation duration, yet behaviourally the ability to discriminate concentration shows no dramatic differences, congruent with the recent findings in rats for a different task (Shusterman et al., 2017). Thus, variable sniffing appears to have no overt negative impact on concentration perception.

### Mitral and tufted cells respond to inhalation changes in cell type specific ways

We have so far shown that it is difficult to distinguish the effect of a change in inhalation or a change in concentration via their effects on MTC responses (Fig. 1–3), yet mice are perfectly capable of fine concentration discrimination in the face of fluctuating inhalations (Fig. 5). We thus conjecture that the olfactory system requires information about the kind of inhalation that just occurred to infer whether concentration or sniffing evoked the response change. This could either be achieved through an efference copy of the sniffing control signal, or reafference (the sensory result of the sniff motor command). Congruent with the latter idea, OSNs have been demonstrated to respond to pressure changes (Grosmaitre et al., 2007), giving rise to sniff coupling in the olfactory bulb (Adrian, 1950; Cang and Isaacson, 2003; Fukunaga et al., 2012; Macrides and Chorover, 1972; Margrie and Schaefer, 2003), which disappears with naris occlusion (Margrie and Schaefer, 2003). We thus wanted to determine whether the olfactory bulb reports changes in single sniffs on a short timescale.

We took baseline activity in absence of odor as a proxy for the large portion of mitral and tufted cells which will not be responsive to an odor, whose activity could instead be used to directly determine the kind of sniff that took place. To do this we analysed the cellular activity of 45 MTCs recorded in passive mice across over 1000-2000 sniffs occurring in absence of the odor. Sniffs were categorised according to inhalation duration, and peristimulus time histograms and average membrane potential waveforms were calculated over 250 ms triggered by inhalation onset for each subset (Fig. 6A-C). We found that individual mitral and tufted cells would show linear transformations in their activity according to the duration of the inhalation just occurring. For example, some cells showed increased spike count (Fig. 6A_1_-B_1_) and depolarising membrane potential (Fig. 6C_1_) as inhalations became faster, while others showed decreasing spike count (Fig. 6A_2_-B_2_) and more hyperpolarising membrane potential (Fig. 6C_2_). 24% of cells showed significant relationships between spike count and inhalation duration (p<0.01, linear regression; Fig. 6D) compared to only 3% in shuffle controls. R^2^ for the actual correlations were also significantly higher than for shuffle controls (actual R^2^ median = 0.54, IQR=0.17-0.82; shuffled median=0.18, IQR=0.04-0.45, p=1×10^−4^, Ranksum, n=41 vs 369; Fig. 6D). Similarly, 22% showed significant correlations with mean membrane potential compared to 2% of shuffle controls (p<0.0, linear regression; Fig. 6D), with R^2^ values for the actual data being significantly higher than for shuffled data (actual R^2^ median = 0.56, IQR=0.20-0.73; shuffled median=0.18, IQR=0.04-0.41, p=9×10^−7^, Ranksum, n=41 vs 369). Timing of activity was also often linearly correlated with inhalation duration, generally with the peak of the membrane potential shifting to earlier times as inhalation duration reduced (significant R values in 32% of cells vs 2% in shuffle controls (Fig. 6D); actual R^2^ median=0.50, IQR=0.22-0.86; shuffled median=0.20, IQR=0.05-0.43, p=2×10^−6^). Altogether 51% of cells showed a significant relationship (p<0.01) between inhalation duration and at least one or more of these activity parameters (Fig. 6E).

**Figure 6.**
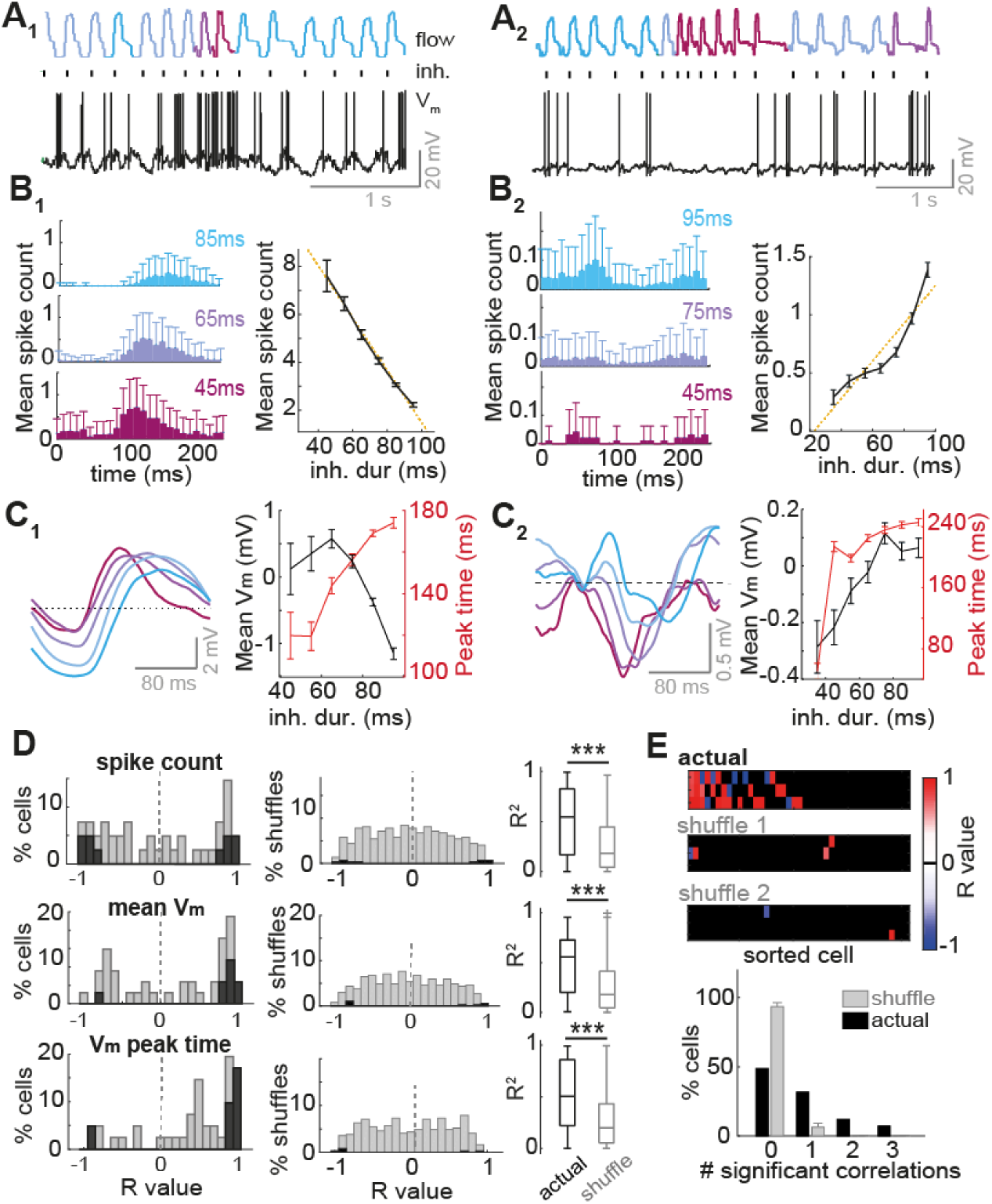
Inhalation duration transforms mean baseline MTC activity in a large proportion of cells. A_1_ to C_1_ refer to one example cell, while A_2_ to C_2_ refer to a different example cell. (**A**) Example nasal flow traces and V_m_ traces in absence of odor. Sniffs are colour coded according to inhalation duration (blue = slow, and red = fast). Black ticks indicate inhalation onset. (**B**) Left: Average spike count triggered by inhalations of different durations (denoted on each histogram). Right: mean spike count per sniff as a function of inhalation duration. Errorbars = standard error (SE). (**C**) Left: Inhalation-triggered mean membrane potential waveforms for sniffs of different inhalation duration. Right: mean V_m_ and timing ofV m peak for membrane potential waveforms across all sniffs as a function of inhalation duration. Errorbars = SE. (**D**) Left: Histograms of R values for correlations between inhalation duration and different activity variables: (from top to bottom) spike count, mean membrane potential, and the timing of peak membrane potential. Middle: histograms of R values between the different activity parameters and inhalation duration for shuffle controls (n = 10 shuffles × 45 cells). Dark bars show significant correlations. Right: Boxplots show comparison of actual R^2^ values for the correlations as compared to shuffle controls. (**E**) Top: Heatmap of R values for correlations between inhalation duration and 3 different activity parameters (spike count, mean membrane potential and timing of peak membrane potential, rows 1-3 respectively), for 45 mitral and tufted cells. Cells are sorted left-right by largest number of significant correlations to lowest number. Black squares show where the correlation was insignificant (p>0.01, regression analysis). Two lowest heatmaps show the same data but for 2 example shuffle controls, where inhalation durations were shuffled with respect to the physiology, and the data re-analysed. This gives an indication of false positive rates in this analysis. Bottom: histogram to show proportion of cells with 0 to 3 significant correlations between the different activity parameters and inhalation duration. Grey shows proportion for shuffle controls.

It has previously been suggested in anaesthetized animals that MCs and TCs are coupled to different phases of the sniff cycle, with MCs coupled to inhalation and TCs to exhalation, allowing designation of putative MC and TC phenotypes based on phase preference (Fukunaga et al., 2012) - something which is supported by recent data from the awake mouse (Jordan et al., 2017). This divergent coupling is thought to be the result of divergent sniff-driven circuit architecture, with TCs largely driven by direct excitation and MC activity heavily modulated in parallel via feed-forward inhibition (Fukunaga et al. 2012). Since reduced inhalation duration caused cells to either depolarise (negative R values between inhalation duration and mean activity – e.g. Fig. 6A_1_-C_1_) or hyperpolarise (positive R values, e.g. Fig. 6A_2_-C_2_), we wanted to test whether this could be explained by cell type. Indeed we found cases where morphologically reconstructed cells showed correlations between certain activity parameters and inhalation duration, with polarity corresponding to cell type as predicted. For example, the tufted cell in Fig. 7A_1_-C_1_ showed significant depolarisation as inhalation duration gets shorter (increased spike count and more depolarised Vm waveform), while the mitral cell in Fig. 7A_2_-D_2_ showed the opposite correlation, showing increasing inhibition as inhalation duration reduced. We plotted R values for these correlations as a function of sniff cycle phase preference (calculated from subthreshold activity) for all cells showing strong correlations (p<0.05, R^2^>0.6). Indeed, for both parameters (mean V_m_ and spike count), there was a significant organisation according to phase for both mean V_m_ (p<0.01, bootstrapping, see methods; Fig. 7D) and spike count (p<0.001; bootstrapping, see methods; Fig. 7E). Moreover, comparing R values between the phase boundaries assigned for putative MCs and TCs resulted in significant differences between the two groups in each case (mean V_m_: pMC: median = 0.93, IQR = 0.84 to 0.95, n = 6; pTC: median = −0.83, IQR = −0.96 to −0.82, n = 10; p = 0.002, Ranksum; spike count: pMC: median = 0.84, IQR = −0.88 to 0.96, n = 22; pTC: median = −0.92, IQR = −0.94 to −0.89, n = 12; p = 0.008, Ranksum).

**Figure 7.**
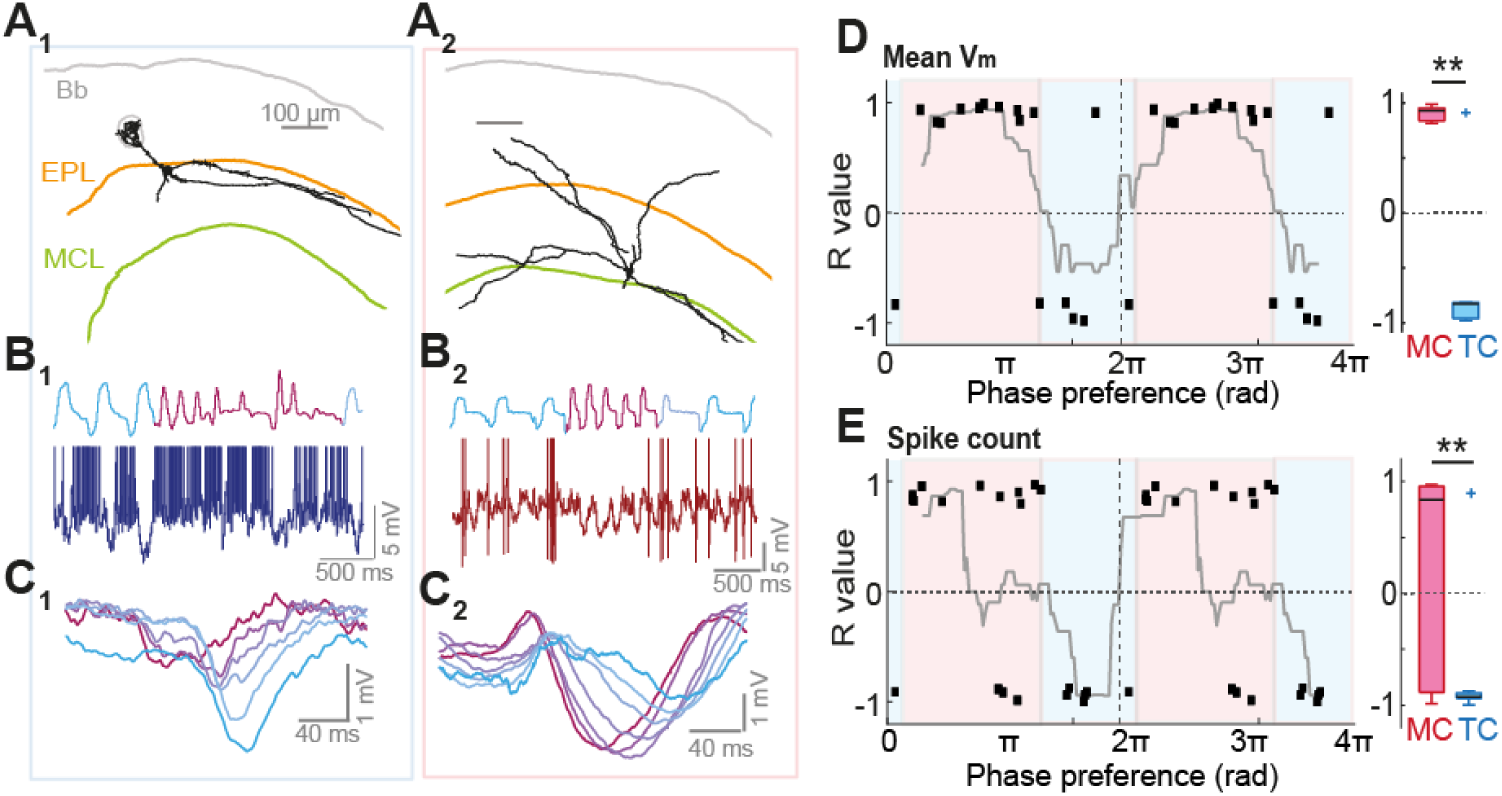
Cell type specificity of effect of inhalation. (**A**_1_) Reconstructed morphology of a tufted cell recorded in awake mouse. ‘Bb’ refers to brain border, ‘EPL’ refers to external plexiform layer and ‘MCL’ refers to mitral cell layer (these morphologies have been previously published in Jordan et al. 2017 for different purposes). (**B**_1_) Example nasal flow and V_m_ trace during a rapid sniff bout (blue to purple represents slow to fast inhalation on flow trace. Spikes have been cropped. (**C**_1_) Mean membrane potential waveform for different bands of inhalation duration: blue = long inhalation duration, purple = short. (**A**_2_-**C**_2_) as for A_1_-C_1_, but for a filled mitral cell recorded in an awake mouse. (**D**) R values for correlations between inhalation duration and mean V_m_ as a function of phase preference. Only strong correlations have been included (p<0.05 and R^2^>0.6). Grey line shows mean R value for all cells within a 2 radian moving window (centred), to give an idea of the phase modulation strength of the data. Boxplots to the right compare all values within the putative MC (red) and putative TC (blue) phase boundaries. (**E**) As for panel D, but for mean spike count per sniff.

Thus, phase locking, which likely relates to MC and TC phenotype, determines how a cell will respond to changing sniff parameters in absence of odor. Thus the large population of cells that are not directly involved with the encoding of odor information could instead be utilised to encode the parameters of each inhalation.

### Inhalation change can be detected and decoded from MTC spiking on rapid timescales

We next sought to determine whether we could read out changes in inhalation from the spiking activity of cells in absence of odor, as a proxy for cells that are not responding directly to the odor.

We first wanted to determine how rapidly a change in inhalation could be detected. For all cells with enough sniff variation (>50 sniffs in each inhalation duration category), we calculated sequences of spike histograms for different inhalation durations using random subsets of sniffs within each group (Fig. 8A; see methods). We constructed either a sequence with PSTHs calculated from three consecutive sniffs of 95 ms inhalation duration, or a sequence with PSTHs calculated from 2 consecutive sniffs of 95 ms, with the last PSTH instead constructed from 55 ms inhalation duration sniffs (Fig. 8A). Using these, it was possible to determine a change in inhalation duration (95 to 55 ms inhalation duration) within only 70 ± 12ms by calculating Euclidean distances between constructed population vectors of the two different sequences (Fig. 8B see methods). Smaller changes in inhalation duration (95 to 75 ms) could also be detected on similarly rapid timescales (Supplementary Fig. 7).

**Figure 8.**
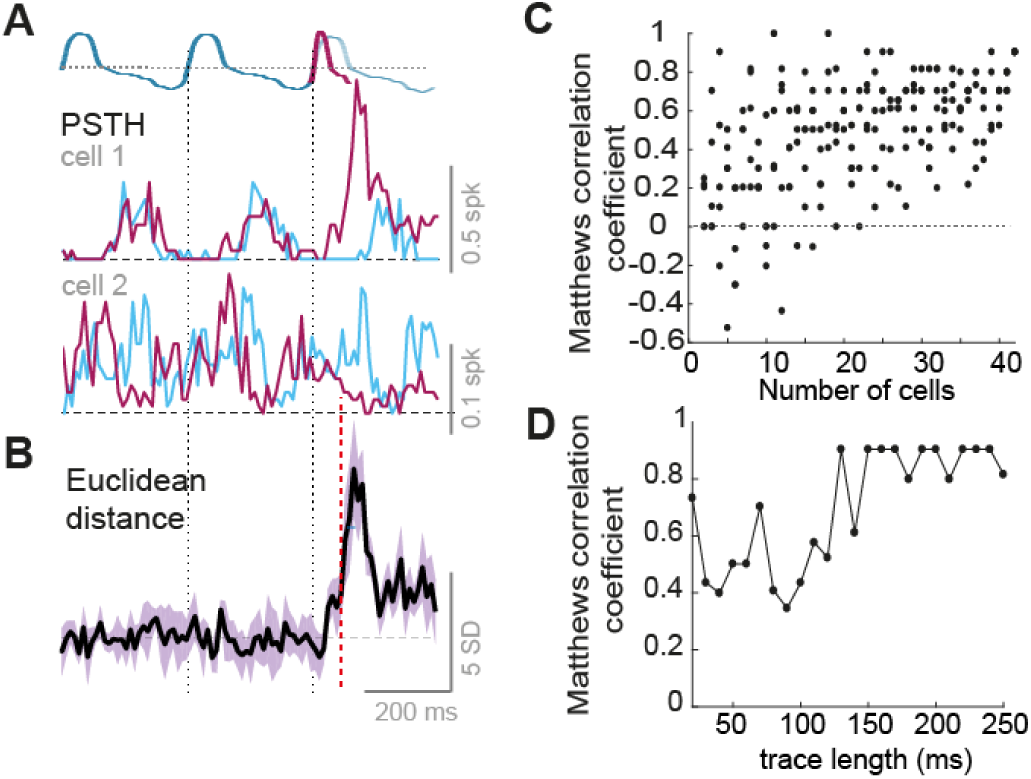
MTC spiking activity is informative of the type of inhalation occurring. (**A**) Top: diagram to show construction of sniff sequences of different inhalation duration: either three of 95 ms inhalation duration, or two of 95 ms with the final sniff of 55 ms duration. Below: two example PSTH sequences averages from random subsets of sniffs that show the particular inhalation duration. Blue line shows the PSTH sequence for 4 sniffs of 95 ms, and purple show sequence in which the last inhalation is of 55ms. (**B**) Mean Euclidean distance calculated between population vectors constructed from the two sniff sequences as in panel A. Plot shows the average of 5 different subsets of data, and shaded area shows standard deviation. Dashed red line indicates time of significant detection of change. (**C**) Classification performance of a trained neural network for distinction of fast and slow inhalations from the emulated population spiking activity constructed from different numbers of MTCs (see methods for details). (**D**) Classification performance for distinction of fast and slow inhalations for the full 42 MTC spiking dataset as a function of the time since inhalation onset.

We next investigated whether the population activity of multiple mitral and tufted cells provides sufficient information for reliable detection of individual sniff cycles with fast and slow inhalation speeds. From 42 single-cell recordings of mitral and tufted cell activity, we constructed an emulated population spiking matrix for respiration cycles for fast (37-80 ms) and slow (96-183 ms) inhalations. After training synaptic weights from all the 42 neurons and an activation threshold, a single output neuron could achieve perfect detection performance of cycles with fast and slow inhalations (Fig. 8C see methods). Reliable discrimination (Matthews correlation coefficient of 0.90) of individual fast and slow inhalation cycles could be achieved within 130 ms after inhalation onset (Fig. 8D).

Thus, even for relatively low numbers of neurons, mitral and tufted cell activity - in absence of odor input - is informative of the inhalation that just occurred, such that non-odor responsive cells could be utilised by the olfactory system to distinguish sniff changes versus concentration changes.

## Discussion

For stable perception, sensory systems must find ways of encoding of stimulus features independent of fluctuating sampling behavior. Here we show that faster sniffs can evoke response changes in the olfactory bulb that appear indistinguishable from those caused by increasing concentration (Fig. 1–3), yet mice are highly capable of perceiving concentration on fast timescales, regardless of sniffing parameters (Fig. 4–5). We reason that the only way the olfactory system can distinguish these two occurrences is via information about the kind of sniff that just occurred. This could potentially be achieved through corollary discharge from a motor circuit involved with breathing rhythms (such as the pre-bötzinger complex). However we find that single MTC activity already correlates with inhalation duration (Fig. 6), and that this is likely generated from feed-forward input in a cell type specific way (Fig. 7), allowing inference about the kind of sniff that just occurred on a rapid timescale (Fig. 8). Thus, the olfactory bulb itself does not appear to be the site where the sniff-invariant percept of intensity is generated, but does appear to already contain the information needed to generate the percept elsewhere.

Given the timescale of decision making for concentration (Fig. 4), it seems likely that the information used by the mouse is the fast timescale temporal shifts in excitation that have been previously described (Cang and Isaacson, 2003; Fukunaga et al., 2012; Sirotin et al., 2015). Congruently, this temporal information contributes to the entirety of concentration discriminability on such a timescale in our dataset (Fig. 3E). It has been suggested that high baseline firing rates of MTCs could obscure such a latency code for concentration being used (Mainland et al., 2014), however this was based on a high estimation of baseline FRs from unit recordings. The whole cell recordings we employ here are thought to be unbiased in terms of baseline FRs (Kollo et al., 2014; Margrie et al., 2002; Shoham et al., 2006), and discriminability of MTC responses based on latency shifts is overt (Fig. 3E). Congruently it is known that mice can perceive the latency difference in optogenetic glomerular activation on the order of tens of milliseconds (Rebello et al., 2014; Smear et al., 2013).

Sniff changes have been hypothesized to alter odor concentration profiles within the nasal cavity (Shusterman et al., 2017; Teghtsoonian et al., 1978). Here we show for the first time directly that sniff changes can indeed mimic the effect of concentration change at the level of both firing rates (Fig. 1), and temporal shifts in spike activity (Fig. 2–3). This is not to say that OSN input is perfectly matched when we compare faster sniff rates and higher concentration. In fact, since subthreshold inhibition is greater for the higher concentration (Supplementary Fig. 3), it would appear that the input strength is higher for the case of increased concentration as compared to faster sniffing. Despite this, overt changes in the spiking output are very similar for increased sniff frequency as compared to increased concentration. Potentially, inhibitory circuits are normalising the spiking output across large changes in input (within a dynamic range), such that while we see differences in subthreshold inhibition, the excitatory spike outputs look very similar. Such a role has been suggested for periglomerular neurons (Roland et al., 2016) and parvalbumin positive interneurons in the external plexiform layer (Kato et al., 2013; Miyamichi et al., 2013).

It has been known for some time that the olfactory bulb is highly modulated by the sniff cycle (Adrian, 1950; Cang and Isaacson, 2003; Fukunaga et al., 2012; Macrides and Chorover, 1972; Margrie and Schaefer, 2003). Since sniff modulation is more overt in anaesthetized mice and is reduced at higher sniff frequencies (Bathellier et al., 2008; Carey and Wachowiak, 2011; Kay and Laurent, 1999), the importance of sniff modulation in the awake animal may come into question. Here we find that sniff patterning of activity gives rise to linear transformations of baseline activity as inhalation parameters are changed, a feature which is widespread throughout MTCs (Fig. 6). We thus reason that a primary function of sniff modulation is to inform the olfactory system of what kind of inhalation took place, such that a change in concentration and a change in sniffing are distinguishable. Congruently we find that inhalation parameters can indeed be readily and rapidly inferred from the spiking activity of MTCs (Fig. 8).

Encoding of ‘sniff effort’ has been hypothesized previously when psychophysics showed that humans could categorise concentrations well despite large changes in inhalation flow rate (Teghtsoonian et al., 1978). Airway resistance is subject to continual changes, and even differs between the two nostrils (Principato and Ozenberger, 1970; Sobel et al., 1999), which will naturally result in varying nasal flow rates for identical respiratory motor commands. Previous work has shown that sniff modulation of the olfactory bulb is generated peripherally rather than centrally, since blocking the naris abolishes sniff modulation in the olfactory bulb (Margrie and Schaefer 2003). Thus reafference using mechanoceptive encoding of sniff pressure, rather than efference copy of the motor commands (which would require constantly updated internal models to calculate the effect on airway flow for each nostril) may be the optimal strategy for encoding inhalation parameters. This could be the reason that olfactory receptors evolved to respond to pressure changes as well as olfactory stimuli (Connelly et al., 2015; Grosmaitre et al., 2007), and indeed may comprise a feature rather than a bug in the olfactory system. Consistently, concentration perception in humans can be affected when the nostril flow rate was changed via experimenter-induced changes in airway resistance instead of volitional changes in sniff pressure (Teghtsoonian and Teghtsoonian, 1984) – i.e. only when flow rate is altered but pressure stays constant. Moreover, imaging of the olfactory cortex in humans identified a region which primarily responds to the sensory effect of sniffing in absence of odor (Sobel et al., 1998).

An accompanying study intuitively suggests that the advance of odor-driven excitation as sniff frequency increases is the result of fluid dynamics in the nasal cavity (Shusterman et al., 2017). A large fraction of our cells show an advance of their baseline activity peak as the inhalation becomes faster (Fig. 6D). We could thus hypothesise that non-odor responsive MTCs within a region of the bulb can provide information about the timing of inspired air reaching the epithelium. If the inhalation becomes faster, both responsive and the much larger population of unresponsive cells show a latency reduction in their peak activity, while if concentration has increased, only the sparse odor responsive population will show this latency shift. Thus, a relative timing code could be used as a sniff-invariant representation of concentration (Supplementary Fig. 9). Exactly where and how the two kinds of information could be integrated to form a sniff invariant representation of concentration should be the objective of future investigations, though recent evidence from the piriform cortex of awake mice already suggests that cortical interneurons sharpen the latency shifts evoked by concentration change and encode concentration via the synchronicity of ensemble firing (Bolding and Franks, 2017). It is also possible that in a mouse performing a concentration guided task, olfactory bulb physiology could be altered by top-down circuits in such a way as to generate a sniff invariant representation of concentration using information about the sniff dynamics.

It is possible that at much larger sample sizes of cells than reported here, a small subpopulation of neurons capable of reporting concentration invariant across sniffs become detectable. It is also possible that we have missed activity parameters at the population level, such as spike synchrony, which could conceivably be more stable reporters of concentration in the face of fluctuating sniffs. We deem these confounds less likely since a complimentary unit recording study finds that the latency shift of excitatory response due to sniff change is widespread throughout a larger sample of MTCs (Shusterman et al., 2017).

In conclusion, concentration changes in the naris can either be self-generated through changes in sniffing, or the consequence of a true change in environmental concentration, yet mice can perform sniff-invariant concentration discrimination. The olfactory bulb contains information about both the odor concentration alongside the inhalation dynamics, which together may allow inference about whether a sniff change or a concentration change occurred, overall enabling sniff-invariant concentration perception.

## Methods

All animal experiments were approved by the local ethics panel of the Francis Crick Institute and UK Home Office under the Animals (Scientific Procedures) Act 1986. All mice used were C57BL/6 Jax males aged between 5 and 12 weeks and were obtained by in-house breeding. All chemicals were obtained from Sigma Aldrich (Missouri, USA).

### Olfactometry

Odorants were delivered to the animal using a custom made olfactometer as used previously (Kollo et al., 2014; Jordan et al., 2017). This consisted of 8 different odor channels connecting two manifolds, a clean air channel, and a final dilution channel carrying clean air. Air was pressure controlled at 1 Mbar with a pressure regulator (IR 1000, SMC Pneumatics, California, USA). Flow was computer controlled to each manifold such that the channel supplying the mouse provided a constant flow of 2000 sccm/N_2_ at all times, meaning that no change in flow accompanied odor pulses. Odor pulses were calibrated to square pulses of different concentrations using a mini photo-ionisation detector (miniPID, Auroras Scientific): briefly, pure odor was presented to the PID from an open bottle, and the maximum recorded voltage (V_max_) was assumed to represent 100% saturated vapor pressure. The pulse amplitudes were then calibrated according to this value, such that a given concentration C (% saturated vapor pressure) could be specified by attaining a square pulse of amplitude equal to C*V_max_/100. Valves and flow controllers were controlled using custom-written LabView software. Odors applied to animals included 2 different odor mixtures (for recordings, either mixture A: Methyl salicylate, eugenol, cinnamaldehyde, creosol and 1-nonanol; or mixture B: guaiacol, valeric acid, (+)-carvone, 2-phenyl ethanol and 4-allylanisol). The components of each mixture were of similar vapor pressure and proportions were adjusted according to relative vapor pressure values as previously described (Jordan et al., 2017). For behavior, either mixture A or pure vanillin odor were applied at various concentrations (Fig. 4 and 5).

### Surgery

Sterile surgical technique was applied during all surgeries. For implantation of the head-plate, mice were anaesthetized with isoflurane in 95% oxygen (5% for induction, 1.5-3% for maintenance). Local (mepivicaine, 0.5% s.c.) and general analgesics (carprofen 5 mg/kg s.c.) were applied immediately at the onset of surgery. An incision was made dorsally above the cranium overlying the cortex and cerebellum, and periosteal tissue was removed. The surface of the bone was drilled away across the implantation surface using a dental drill, and cyanoacrylate was applied to the sutures between the cranial bones to reduce movement. A stainless steel custom head-plate was then glued to the bone surface with cyanoacrylate, and dental cement was used to reinforce the bond. For mice going on to whole cell recording, an additional chamber was constructed on the bone overlying the right olfactory bulb using dental cement. After surgery, the mice were allowed to recover for 48 hours with access to wet diet.

### Whole cell recordings

On the day of recording, mice were again anaesthetized with isoflurane as above, and carprofen analgesic injected (5 mg/kg s.c.). A 1 mm craniotomy was made overlying the right olfactory bulb, and the dura removed. A layer of 4% low melting point agar was then applied to the surface of the bulb, about 0.5-1 mm deep to reduce brain movement. Cortex buffer (125 mM NaCl, 5 mM KCl, 10 mM HEPES, 2 mM MgSO_4_, 2 mM CaCl_2_, 10 mM glucose) was used to fill the recording chamber. The animal would then be transferred to the recording rig, and allowed to awake from anaesthesia for 20 minutes. Whole-cell recordings were then made blindly by descending a 5-7 MΩ borosilicate glass micropipette (Hilgenberg, Malsfeld, Germany) filled with intracellular solution (130 mM KMeSO_4_, 10 mM HEPES, 7 mM KCl, 2 mM ATP-Na, 2 mM ATP-Mg, 0.5 mM GTP, 0.05 mM EGTA, and in some cases 10 mM biocytin; pH adjusted to 7.4 with KOH, osmolarity=280 mOsm) through the agar and 180 μm into the olfactory bulb with high pressure. Here pressure was reduced, and the micropipette advanced in steps of 2 μm until a substantial and sudden increase in resistance was observed indicating proximity to a cell. Pressure was then dropped to zero or below, and a gigaohm seal attained. Whole-cell configuration was then achieved, and the membrane voltage recording made in current clamp mode. Identification of mitral and tufted cells was made using electrophysiological parameters, such as in input resistance below 150 MΩ, a resting membrane potential between −60 and −40 mV, and an after-hyperpolarisation waveform conforming to MTC phenotype in an independent component analysis performed as previously (Kollo et al., 2014; Jordan et al., 2017).

Altogether 14 cells were recorded in passive mice and presented with 2 different odor concentrations, as well as puff stimuli to evoke fast sniffing (Fig. 1 and 3). Some cells were presented two different odor stimuli (two different mixtures), such that there were 20 cell-odor pairs in total. Concentrations were presented in a pseudorandom order and puff stimuli occurred on a random subset of trials only for the low concentration. Puff stimuli were applied simultaneous with the odor stimuli with a gentle clean air stream to the flank. For some analyses, e.g. Fig. 2 and for Fig. 6–8, data was supplemented with previously recorded cells from the passive mouse (Jordan et al., 2017) presented the same odor mixtures at 1% vapor pressure (n = 6 and n = 38 respectively).

### Behavioral task and training

On day 0 (48 hours after surgery), mice with head-plates implanted would begin water restriction. On day 1, mice were habituated to the experimenter and hand-fed 0.5 ml of highly diluted sweetened condensed milk with a Pasteur pipette. On day 2, mice were habituated to head-fixation: mice were head-fixed above a treadmill and allowed access to free reward upon licking. On day 3, successfully habituated mice underwent operant conditioning with repeated presentations of CS+ concentration of the odor mixture until the mouse learned to lick in the 1 s after odor offset to receive the reward. On day 5, the CS- concentration was also presented alongside the CS+ concentration in a pseudorandom order, until the mice learned to refrain from licking to the CS-. Licking to the CS-would evoke an addition of 6 s to the inter trial interval. 5 mice were trained with high concentration stimuli as the CS+ (‘high go’), and 3 mice were trained on the reverse contingency (‘low go’). On day 6-8, mice would be presented with 5 different concentrations (3 additional concentrations spanning the range between the previously two learned concentrations), and contingencies as depicted in Fig. 5A. On day 9, five mice went on to a final session: after observing criterion performance on the binary odor concentration task with the mixture as learned previously, the odor would switch to vanillin with the same contingency between concentrations. Mice were carefully monitored to maintain their body-weights above 80% of their pre-restriction weight and were ensured a minimum of 1 ml water per day regardless of performance. Any mouse showing signs of distress were immediately returned to water access.

### Data analysis

#### Statistics

In all cases, 5-95% confidence intervals were used to determine significance unless otherwise stated. In all figures, a single asterisk denotes p<0.05, double asterisk denotes p<0.01 and a triple asterisk denotes p<0.001. Means and error bars showing a single standard deviation either side are used in all cases for normally distributed data of equal variance. Two-sided Student’s t-tests were used for comparison of means and Bartlett tests used to compare variances, unless otherwise stated. Boxplots are used to represent any other data (data comparisons of unequal variance, or non-normally distributed data), where median is plotted as a line within a box formed from 25th (q1) and 75th (q3) percentile. Points are drawn as outliers if they are larger than q3 + 1.5 × (q3 - q1) or smaller than q1 – 1.5 × (q3 - q1). For such data, Ranksum tests were used to compare the medians, and Browne-Forsythe tests used to compare variance, unless otherwise stated.

#### Sniff parameters

Using the recording of nasal flow, different sniff parameters could be extracted. First, inhalation peaks were detected using Spike2 algorithms that mark each peak above a certain threshold voltage manually defined by the user, such that all inhalations were included and no false positives were present. Inhalation onset was defined as the nearest time-point prior to inhalation peak at which the flow trace reached zero. Inhalation offset was similarly calculated as the first time point after inhalation peak where the flow trace reached zero. Inhalation duration was defined as the difference in time between inhalation onset and offset. Peak inhalation slopes were calculated by detecting the peak value of the flow waveform differential 50-0 ms prior to inhalation peak. Sniff duration was calculated as the time between subsequent inhalation onsets. Sniff frequency was calculated by taking the inverse of the mean sniff duration within the odor time period.

#### Spike rate responses and onsets

Long timescale (Fig. 1): for each cell, mean spike count was calculated in 250 ms time bins for the full 2 s odor stimulus. These were then averaged across trials to generate PSTHs for low concentration and fast sniffing (5 trials of lowest mean inhalation duration), low concentration and slow sniffing (5 trials with highest mean inhalation duration) and high concentration and slow sniffing. Values were quadrupled to estimate FR in Hz.

Short timescale (Fig. 2–3): for each cell, spike counts were calculated in 10 ms time bins for only the first 250 ms from odor onset (aligned to first inhalation onset). These spike counts were then averaged across trials for low concentration and fast inhalation (>70^th^ percentile peak inhalation slope), low concentration and slow inhalation (<30^th^ percentile inhalation slope) and high concentration and slow inhalation (<30^th^ percentile). Onset for excitatory responses was defined at the point the mean spike count exceeded the mean + 2 SDs of the baseline spike rate in the 250 ms prior to odor onset, and remained there for at least 2 consecutive points.

#### V_m_ responses

To analyse subthreshold responses in absence of spiking activity, spikes and their AHPs were subtracted from the trace. This was done by first using the ‘wavemark’ tool in Spike2 to detect spikes by thresholding and matching them to a generated spike waveform template. The length of this spike waveform template was manually adjusted for each cell according to its AHP length, but was usually around −4ms to 20-30 ms post spike peak. A trace was then generated containing all detected spike waveforms connected by zero values, and this was subtracted from the original voltage trace.

#### Correlations between response changes due to sniffing and concentration change

For both long and short timescale mean FR responses, changes in FR response were calculated for sniff change (fast-slow sniffing, low concentration odor) and concentration change (high-low concentration, slow sniffing). For all cell-odor pairs across the sample, a single regression was made between FR changes for sniff change and FR changes for concentration change in the corresponding time bins, generating an actual R and p value (Fig. 1E and 3C). For shuffle controls, low concentration trials were shuffled in respect to the sniff behaviour on each trial, and the same analysis was repeated 100 times.

#### Euclidean distance analysis of concentration discriminabilty

In reference to Fig. 3E and Supplementary Fig. 5G: Euclidean distance was taken across the population between mean spike counts for high concentration and low concentration (slow inhalation). This generated a measure of discriminability between concentrations when the inhalation was slow for both concentrations. To test how much of the discriminability was due to latency shift of excitation, responses which underwent a detectable latency shift between high and low concentrations had their spike count response to low concentration manually shifted forward according to the latency shift occurring between high and low concentration. Euclidean distance was then recalculated between spike counts for high concentration and the latency-shifted spike counts at low concentration. Finally, Euclidean distances were taken between spike counts for high concentration (slow inhalation) and low concentration (fast inhalation). Time for discrimination was calculated, if possible, as the point at which the Euclidean distance exceeded the mean + 2 SDs of the baseline Euclidean distance (250 ms prior to odor onset) for at least 2 consecutive 10 ms time bins.

#### Baseline activity correlations with inhalation duration

For each cell (n = 48), 1000-2000 sniffs were analysed in absence of odor. Sniffs were categorised according to their inhalation duration, 35-45 ms, 45-55 ms, 55-65 ms and so forth. For each individual sniff, different parameters were calculated from the corresponding neural activity. Mean membrane potential was calculated from the subthreshold membrane potential occurring from 0 to 250 ms from inhalation onset. Peak membrane potential was designated as the maximum membrane potential within 30-250ms after inhalation onset, and time of the peak membrane potential was calculated at the time of this maximum membrane potential relative to inhalation onset. Spike counts were calculated by summing all action potentials occurring within the same timeframe. To calculate the correlations for each parameter, each was averaged across all sniffs within the category and regression analysis was used to generate an R and p value between the resulting average parameters and the corresponding inhalation duration (minimum of the category). For each cell, inhalation duration categories were excluded from the correlation if they contained less than 25 sniffs, and cells that had less than 5 valid categories were additionally excluded. For shuffle controls, inhalation duration was shuffled throughout the data and the regression analysis repeated 10 times per cell.

#### Euclidean distance analysis of detectability of sniff change

For this analysis only cells with more than 50 sniffs during baseline in each category: 55-65 ms, 75-85 ms and 95-105 ms inhalation duration were included. A random subset of 25 sniffs in each group were selected and spike activity within these samples were used to construct PSTHs. PSTHs were put in sequence, either 3 consecutive 95 ms inhalation duration sniffs (control sequence), or the same sequence but with the final sniff of a different inhalation duration, either 75 ms or 55 ms. Each PSTH was normalised such that the first 30 ms started at zero Hz on average. Euclidean distance across the population of these sequences were then calculated between the control sequence and sequences ending in 55 ms or 75 ms inhalation duration sniffs. Detection time for the change in inhalation duration was calculated where the Euclidean distance in the last sniff exceeded the mean + 2 SDs of the baseline Euclidean distance from the first 2 sniffs.

#### Discriminability of slow and fast sniffs from emulated MC/TC population activity

Sniff cycles from 42 recorded neurons (25 MC, 17 TC) were divided into fast (37-80 ms), medium (80-96 ms) and slow (96-183 ms) cohorts. All cycles with sniff duration below the 0.5th percentile (108 ms) and above the 99.5th percentile (597 ms) were discarded. All cycles with inhalation duration below the 0.5th percentile (37 ms) and above the 99.5th percentile (183 ms) were discarded. From each cell the spiking activity for 400 ms (except for Fig. 8D, where the length was varied) following the inhalation onset in a randomly chosen fast or slow inhalation cycle was combined to create emulated population spike trains (32 slow and 32 fast inhalation cycles). The emulated inhalation cycles split to a training set (22 slow and 22 fast inhalation cycles) and a test set (10 slow and 10 fast inhalation cycles). For each MC/TC neuron a postsynaptic EPSPs waveform (one sample per ms) was simulated by convolving the spike train with a normalised alpha function (amplitude=10, time constant: 10 ms, see Kollo et al. 2014). The compound EPSP of the read-out neuron was calculated as the weighted linear sum of the individual EPSP waveforms of each cell (Supplementary Fig. 8). The read-out neuron was activated if the maximum of the EPSP waveform crossed the threshold value. To find the optimal values of EPSP weights and threshold, the Nelder-Mead method was used with a logistic activation function in the read-out neuron. Discrimination performance of fast and slow inhalation cycles was assessed on the test set with the Matthews correlation coefficient.

#### Phase preference and putative MC and TC boundaries

The theta modulation properties of each cell were calculated as previously (Fukunaga et al. 2012; Jordan et al., 2017). Due to the high variability of sniff behaviour in awake mice, analysis was restricted to sniff cycles between 0.25 and 0.3 s in duration, where the preceding sniff cycle was also within this range. Mean V_m_ from the spike-subtracted Vm trace was taken as a function of sniff cycle phase for at least 150 sniffs, and this was plotted as Cartesian coordinates. The angle of the mean vector calculated by averaging these Cartesian coordinates was taken as the phase preference of the cell. To determine putative MC or TC type based on phase preference, we used the phase boundaries calculated as previously (Jordan et al. 2017).

#### Modulation of sniff-activity relationships across phase preference

In reference to Fig. 7D and E. To determine whether the sign of relationships between inhalation duration and the various activity parameters is modulated by the sniff phase preference of the cell, R values for the various correlations were plotted as a function of phase preference. Only correlations with a significant p value (<0.05) and with an R^2^>0.6 were included. A sliding window of 2 radians was then used to calculate the mean R value for all cells with phase preference within the window, resulting in a mean R value as a function of phase preference. The modulation strength of mean R value as a function of phase was then calculated: the plot of mean R value was normalised to the minimum value across all phases, and the result plotted as Cartesian coordinates. The length of the mean vector calculated by averaging these Cartesian coordinates was taken as the modulation strength of the R value across phase space. To determine the significance of this modulation, R values were shuffled with respect to phase preference 10000 times, and the resulting distribution of shuffled modulation strength was compared to the value for the unshuffled data.

#### Learning time and reaction time

##### Learning time

For the generation of learning curves as in Fig. 4, a moving window was used across 5 consecutive CS+ and 5 consecutive CS-trials, advanced by 1 trial on each step, and a percentage correct calculated. The trial at which this reached 80% correct for 5 consecutive points was deemed the learning time.

Reaction time calculations were based on 10 or more trials of 80% performance. **From lick behavior:** For each CS+ and CS-, lick probability was calculated in a moving time window of 100 ms, aligned to the first inhalation after final valve opening. The difference between the probability of licking for CS+ and CS-for each time window was calculated, and the leading edge of the first window at which this calculated difference significantly deviated from the values calculated from the 2 s window prior to odor onset was considered the reaction time. **From sniff behavior:** inhalation and exhalation duration values were calculated for CS+ and CS-as a function of sniff number from odor onset. These values were compared between those calculated for CS+ and CS-using a t-test, and the reaction time was calculated based on the first inhalation or exhalation within the series to show a significant difference.

## Author Contributions

A.T.S., and R.J., designed all experiments, R.J. performed all experiments, and analysed all data apart from the discriminability of fast and slow sniffs (Fig. 8C-D), which was performed by M.K. The article was written by R.J. and M.K. with contributions from A.T.S. The authors declare no competing financial interests.

## Acknowledgments

We thank Martyn Stopps and Nicholas Burczyk for assistance with custom made equipment, Mostafa Nashaat and Edward Bracey for initial support with behavioral training, Christoph Schmidt-Hieber for advice on whole-cell recording *in vivo,* and Roma Shusterman, Andrew Erskine, Christina Marin, Izumi Fukunaga and Kevin Bolding for helpful comments on the manuscript. This work was supported by the Francis Crick Institute which receives its core funding from Cancer Research UK (FC001153), the UK Medical Research Council (FC001153), and the Wellcome Trust (FC001153); by the UK Medical Research Council (grant reference MC_UP_1202/5); by the DFG (SPP 1392); and a Boehringer Ingelheim Fonds PhD fellowship to RJ. AS is a Wellcome Trust Investigator (110174/Z/15/Z).

